# Chromosome-wide evolution and sex determination in a nematode with three sexes

**DOI:** 10.1101/466961

**Authors:** Sophie Tandonnet, Georgios D. Koutsovoulos, Sally Adams, Delphine Cloarec, Manish Parihar, Mark L. Blaxter, Andre Pires-daSilva

**Affiliations:** School of Life Sciences, University of Warwick, Coventry CV4 7AL, UK; Institute of Evolutionary Biology, University of Edinburgh, Edinburgh EH9 3JT, UK; INRA, UMR1355 Institute Sophia Agrobiotech, F-06903 Sophia Antipolis, France; Department of Molecular Medicine, Institute of Biotechnology, University of Texas Health Science Center at San Antonio, San Antonio, TX, USA

## Abstract

The free-living nematode *Auanema rhodensis* is a rare example of a species with three sexes, in which males, females and hermaphrodites coexist. *A. rhodensis* males have only one X chromosome (XO), whereas females and hermaphrodites have two (XX). The *A. rhodensis* X chromosome is unusual: it does not recombine in hermaphrodites and is transmitted from father to son. The mechanism that controls the production of females *versus* hermaphrodites is unknown but is dependent on maternal and larval environmental factors. Here we report the genome sequence and genetic map of *A. rhodensis*, placing over 95% of the sequence in seven linkage groups. Comparison of the seven *A. rhodensis* chromosomes to chromosomal assemblies of *Caenorhabditis elegans* and three other rhabditine nematodes identified deeply conserved linkage groups we call Nigon units, some of which have been maintained in all species analysed. Others have undergone breakage and fusion, and the *A. rhodensis* karyotype is the product of a unique set of rearrangements involving three Nigon units. The *A. rhodensis* X chromosome is much smaller than the autosomes, is less gene dense and is 3 to 4 times more polymorphic, reflecting its unique transmission history. Differential expression analyses comparing females and hermaphrodites, identified several candidate genes, including orthologues of *C. elegans gld-1*, *tra-1* and *dmd-10/11*, that are potentially involved in the female-hermaphrodite sexual decision.

## Introduction

Multicellular organisms exhibit a wide spectrum of breeding systems. The most common include gonochorism (male/female), hermaphroditism (self-compatible or not) and parthenogenesis. In addition to these, there are rare mixed breeding systems, which include androdioecy (males and hermaphrodites), gynodioecy (females and hermaphrodites) and trioecy (co-occurrence of males, females and hermaphrodites) [1]. Mating strategies heavily influence the genetics and consequently the evolution of populations [2]. For example, because of the absence of or ineffective recombination, selfing is predicted to reduce the effective population size, decrease the rate of appearance of novel allelic combinations and increase linkage disequilibrium and genetic hitchhiking, and thus lead to an overall reduction in the level of polymorphism [2-4]. On a macro-evolutionary scale, selfing can cause isolation between populations and result in local extinction, since low effective population size combined with low genetic diversity decreases the efficacy of selection to remove deleterious alleles and adapt to new environments [2-4]. However, under certain conditions (e.g. low population density) selfing offers reproductive assurance, as self-compatible individuals can reproduce without a mating partner. Outcrossing (gonochorism, self-incompatible hermaphroditism) favours genetic diversity and offers a greater potential to adapt to changing environments at the cost of the necessity of finding a mate. The costs and advantages of outcrossing *versus* selfing depend on environmental factors and thus selection may favour transitions between mating systems. Androdioecy, gynodioecy and trioecy are usually thought to be intermediate evolutionarily unstable strategies [1], but offer important systems to test models of the causes and consequences of mating system on the evolution of populations.

Species in phylum Nematoda (roundworms) exhibit a wide range of mating systems and are thus an ideal group for understanding the origin and maintenance of alternative mating strategies. Hermaphroditism and parthenogenesis have evolved many times independently in Nematoda, in most cases most probably from ancestral gonochoristic ancestors [5, 6]. Mixed breeding strategies, including trioecy and heterogony (alternating generations of distinct mating strategies) are also present [7, 8] The best-known nematodes with mixed breeding strategies are parasites. For example, *Strongyloides* species (Tylenchina, Clade IV [9]) have a facultative free-living generation with gonochoristic females and males, and a parthenogenetic female generation that lives inside the mammalian host [10, 11]. *Heterorhabditis* nematodes (Rhabditina, Clade V) are insect parasites in which hermaphrodites initially produce males, females and hermaphrodites when inside the host, and then shift to the production of infective larvae that are fated to become hermaphrodites [12, 13]. *Rhabdias* species (Rhabditina, Clade V) show strict alternation between a parasitic generation with selfing hermaphrodites, and free-living generation with gonochoristic males and females [14]. Understanding the evolution of these complex life cycles is hampered by the close association with parasitism, and the experimental difficulties that this brings.

Free-living trioecious nematodes, such as *Auanema rhodensis* and *Auanema freiburgensis* [7, 15] are much more experimentally and genetically tractable [16], and may permit delineation of the mechanisms of sex determination and of the processes that underlie the origin and maintenance of alternative mating systems. In *Auanema*, females and hermaphrodites are karyotypically identical, with two X chromosomes (XX), whereas males have only one X chromosome (XO)[17]. Under normal conditions, XX larvae fated to become hermaphrodites obligately pass through a dauer larval stage (L3d), whereas XX larvae fated to be females and XO male larvae do not [7, 15, 18]. The mechanism controlling the female *versus* hermaphrodite fate is unknown, although the sex and age of the mother affect the proportion of XX females *versus* XX hermaphrodites in the progeny of *A. rhodensis* [19].

In *Auanema rhodensis*, females and hermaphrodites can be distinguished in early first stage larvae (L1) by the size of the gonad primordia [15, 18]. Larvae fated to become hermaphrodites have smaller gonad primordia than larvae that develop into females. In *Caenorhabditis elegans* and other rhabditine nematodes, dafachronic acid (DA), a steroid hormone, prevents dauer entry by binding to the nuclear steroid receptor DAF-12. Treatment of hermaphrodite-fated *A. rhodensis* larvae (L1 with smaller gonad primordia) with exogenous DA converts them into females (dubbed ‘converted females’) [18]. Conversely, treatment with dafadine, an inhibitor of the cytochrome P450 DAF-9 that is needed for DA production, converts female-fated *A. rhodensis* larvae (L1 with larger gonad primordia) into hermaphrodites by promoting dauer entry.

In *A. rhodensis* the meiotic program governing the segregation of the X chromosome is determined by the sex of the individual and the type of gametogenesis [20, 21]. The X chromosome does not recombine during meiosis in the germline of hermaphrodites, which produce sperm with two X chromosomes (diplo-X sperm) and oocytes with no X (nullo-X oocytes). Behaviour of the X in females is largely normal, producing mostly recombined haplo-X oocytes, though nullo-X oocytes are produced infrequently. The X is transmitted directly from father to son during cross-fertilization, as males produce exclusively haplo-X sperm (the nullo-X counterpart is provided by the XX individuals) [17, 20, 21]. The biological significance of these meiotic variations and the unusual inheritance of the X are not fully understood, but predict that the *A. rhodensis* X chromosome will have very distinct population genetic characteristics compared to the autosomes, and compared to X chromosomes in other XX-XO nematode species. *A. rhodensis* also has an unusual karyotype compared to most other rhabditine nematodes, with six autosomes in addition to the X [20]. How this karyotype has arisen from the ancestral five autosomes plus X karyotype, observed in most Rhabditina, is unknown.

Understanding the logic of exceptions to biological rules can lead to a greater understanding of the forces that maintain the norm. To explore the mechanisms and evolution of trioecy in *A. rhodensis*, we have developed a high-contiguity genome and genetic map and used this to explore the origins of the X chromosome and its differences from the autosomes. We also compared gene expression in early second-stage larvae (L2) of normal females, converted females and hermaphrodites to identify genes that may play roles in sex determination.

## Materials and Methods

### Strains

We used *A. rhodensis* inbred strains APS4 and APS6 [7] to produce Advanced Intercross Lines (AILs) and generate a genetic linkage map. Strains were maintained at 20 ° C according to standard conditions as for *C. elegans* [22] on Nematode Growth Medium (3 g/L sodium chloride, 2.5 g/L bacto peptone, 17 g/L agar, 1 mM magnesium sulfate, 5 mg/L cholesterol, 1 mM calcium chloride, 25 mM potassium phosphate) [23]. Plates were seeded with the *Escherichia coli* streptomycin-resistant strain OP50-1. Microbial contamination was prevented by adding 50 µg/mL of streptomycin and 10 µg/mL of nystatin to the NGM.

### DNA extraction, sequencing and data pre-processing

To extract nematode DNA with minimal bacterial contamination, we used the split plate method [24]. Nematodes were cultured on one compartment of a 10 cm, two-compartment plate. The compartment with nematodes contained NGM seeded with *E. coli* OP50-1, and the second compartment contained M9 buffer. As the compartment with nematodes became crowded, dauer larvae migrated to the compartment with M9. The dauers were collected from 10 plates and washed twice with M9 buffer. The nematode pellet was stored at −80 °C. DNA was extracted and treated with RNAse using the Gentra Puregene Core A Kit (Qiagen) following the manufacturer’s instructions. The DNA was dissolved in nuclease-free water for library preparation and sequencing.

The APS4 strain was chosen for genome assembly. Three independent Illumina paired-end (PE) libraries with insert sizes of 250 bp were sequenced at UT Southwestern (Dallas, Texas, USA) on an Illumina HiSeq 2500 (Table 1). Two Illumina mate-pair (MP) libraries with virtual insert sizes of ~3 kb and ~6 kb were constructed and sequenced on Illumina HiSeq 2500 at Edinburgh Genomics (University of Edinburgh, UK). The raw reads were processed to remove low-quality bases using Skewer (version 0.2.1, parameter settings “-Q 20 −l 51 −t 32”) [25]. Error correction was performed using Fiona (version 0.2.1, “-nt 48 −g 60000000”) [26]. We used Blobtools [27] to remove microbial contamination.

**Table 1.**
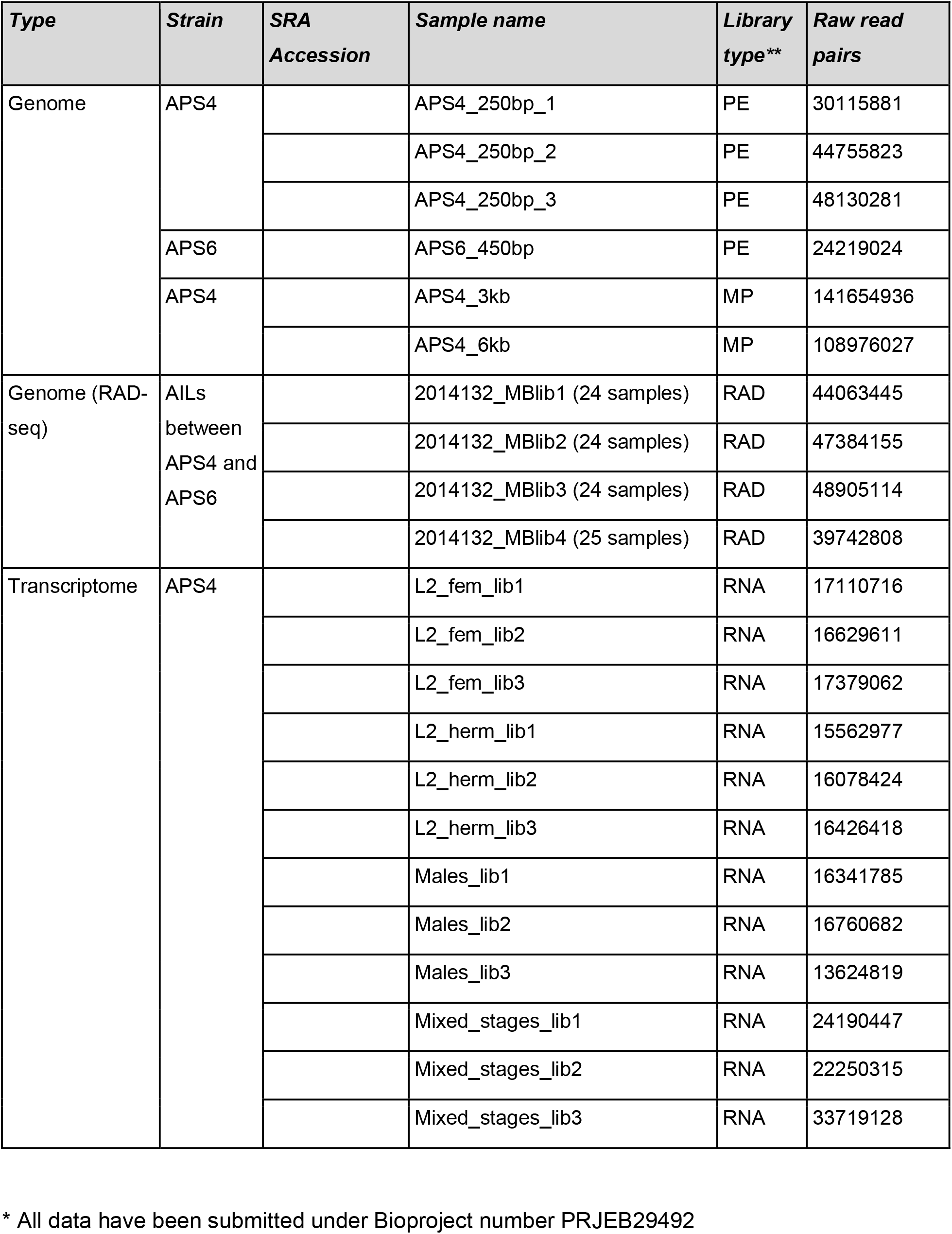
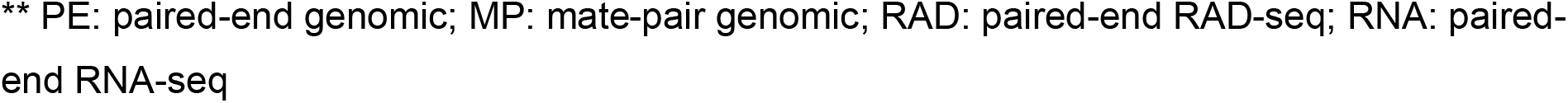
Genome and transcriptome libraries used in this study*.

| *Type* | *Strain* | *SRA*<br>*Accession* | *Sample name* | *Library*<br>*type\*\** | *Raw read*<br>*pairs* |
| --- | --- | --- | --- | --- | --- |
| Genome | APS4 |  | APS4_250bp_1 | PE | 30115881 |
|  |  |  | APS4_250bp_2 | PE | 44755823 |
|  |  |  | APS4_250bp_3 | PE | 48130281 |
|  | APS6 |  | APS6_450bp | PE | 24219024 |
|  | APS4 |  | APS4_3kb | MP | 141654936 |
|  |  |  | APS4_6kb | MP | 108976027 |
| Genome (RAD-seq) | AILs<br>between<br>APS4 and<br>APS6 |  | 2014132_MBlib1 (24 samples) | RAD | 44063445 |
|  |  |  | 2014132_MBlib2 (24 samples) | RAD | 47384155 |
|  |  |  | 2014132_MBlib3 (24 samples) | RAD | 48905114 |
|  |  |  | 2014132_MBlib4 (25 samples) | RAD | 39742808 |
| Transcriptome | APS4 |  | L2_fem_lib1 | RNA | 17110716 |
|  |  |  | L2_fem_lib2 | RNA | 16629611 |
|  |  |  | L2_fem_lib3 | RNA | 17379062 |
|  |  |  | L2_herm_lib1 | RNA | 15562977 |
|  |  |  | L2_herm_lib2 | RNA | 16078424 |
|  |  |  | L2_herm_lib3 | RNA | 16426418 |
|  |  |  | Males_lib1 | RNA | 16341785 |
|  |  |  | Males_lib2 | RNA | 16760682 |
|  |  |  | Males_lib3 | RNA | 13624819 |
|  |  |  | Mixed_stages_lib1 | RNA | 24190447 |
|  |  |  | Mixed_stages_lib2 | RNA | 22250315 |
|  |  |  | Mixed_stages_lib3 | RNA | 33719128 |

\* All data have been submitted under Bioproject number PRJEB29492
\*\* PE: paired-end genomic; MP: mate-pair genomic; RAD: paired-end RAD-seq; RNA: paired-end RNA-seq

To call variants in strain APS6, an Illumina paired-end library with insert size 450 bp was constructed and sequenced on Illumina HiSeq2500 at UT Southwestern (Table 1). Raw reads were preprocessed using Skewer as for the APS4 libraries [25].

### Genome assembly and annotation

*De novo* genome assembly was performed with SOAPdenovo2 (version r240) [28], k-mer length=71) using the PE and MP libraries for contig assembly and MP libraries for scaffolding, as this resulted in the best assembly. The optimal k-mer length was estimated using Kmergenie (version 1.6741) [29]. We removed small contigs (<500 bp) or those having a low coverage (<5 reads/scaffold on average). Reapr (version 1.0.17) [30] was used to identify mis-assemblies within the scaffolds, and 42 questionable joins in the draft scaffolds were identified. We manually inspected these using Tablet (version 1.14.11.07) [31] and manually split 4 scaffolds that contained unjustified joins. Repeats were masked with RepeatModeler [32] and RepeatMasker [33].

Genome completeness was assessed using CEGMA (version 2.4) [34]. Gene predictions were made using the *ab initio* and evidence-driven gene predictors GeneMark (version 2.3)[35], SNAP (trained with CEGMA, release 11/29/2013) [36], Maker2 (version 2.31) [37] and Augustus (version 2.5)[38]. The outputs from SNAP, GeneMark, a set of *A. rhodensis* transcripts assembled by Trinity [39] and protein similarity matches from the UniProt database were used as inputs for Maker2. We then used the output from Maker2 along with hints directly generated using RNA-seq reads and the set of transcripts as input for Augustus. We used the Augustus gene predictions for our analyses.

We functionally annotated the protein coding genes by combining the results from BLAST, InterProScan and Blast2GO. We performed a similarity search (-evalue 1e-5-max_target_seqs 50 -outfmt 5) against a database of all metazoan protein sequences available on NCBI (28/08/2015) using BLAST+ (version 2.2.31+) [40]. InterProScan (-goterms -iprlookup, version 5.14-53.0) [41] was used to identify protein motifs and signatures. We used Blast2GO (version 3.1) [42] to integrate the InterProScan and BLAST results to add Gene Ontology (GO) term annotations to *A. rhodensis* proteins. We added implicit GO terms to the existing annotation using Annex [43].

We identified non-coding RNA loci using Infernal (version 1.1.1) [44], which uses the Rfam database. Transfer RNAs were identified using RNAscanSE (version 1.3.1) [45]. We identified ribosomal RNAs with Infernal (for 5S rRNAs) and BLASTn (BLAST+, version 2.2.31+ [40]) using as a database the partial 18S (accession number EU196004.1) and 28S (accession number EU195960.1) of *A. rhodensis* [46]. We counted unique functional RNA features using BEDTools intersect (version 2.25.0, −s −c, [47]).

### RAD-seq and Genetic Map Construction

A genetic map was constructed using markers obtained from restriction site-associated DNA sequencing (RAD-seq) of 95 advanced intercross lines (AILs) [48] derived from *A. rhodensis* inbred strains APS4 and APS6 [7]. To generate the AILs, crosses between APS4 females and APS6 males were performed to generate several F1 hermaphrodites, which were allowed to reproduce by selfing. Each progeny AIL was established from single F2 hermaphrodite progenitors, which were left to expand. We used the split plate method as described above to isolate DNA from the lines. This method relies on the isolation of dauers. Since dauers of *A. rhodensis* always develop into hermaphrodites [15, 18], the DNA isolation was derived only from this sex. Paired-end RAD-seq using PstI restriction digestion was carried out for each of the parental strains and the 95 AILs [49]. The raw RAD-seq reads were demultiplexed and low-quality regions were removed using process_RAD_tags from the Stacks package (version 1.35) [50]. We then used the denovo_map.pl Stacks pipeline to determine the genotype of each locus (region sequenced adjacent to the PstI cut site) for each progeny sample.

The genetic map was constructed using the R packages OneMap (version 2.0-4)[51] and r/qtl (version 1.38-4) [52]. A LOD (logarithm of odds) score of 20 and a recombination fraction of 0.5 were used as parameters to arrange the loci into linkage groups. The initial genetic map was refined by removing duplicated markers (markers with exactly the same genotype across all samples) and those with missing genotypes in 50% or more of the samples (function ‘drop.markers’). Large gaps (loose markers) in the genetic map were fixed by dropping 3 markers. The Kosambi mapping function was used to determine the genetic distances between markers. However, the genetic distances could not be estimated precisely, as the level of recombination in the AILs is unknown.

### Synteny analysis and identification of the X chromosome

The software Chromonomer (version 1.07) [53] was used to anchor the genomic scaffolds to the genetic map, yielding a chromosomal assembly with scaffolds ordered, where possible, in each linkage group. The resulting chromosomal blocks were aligned to the *C. elegans* and *Pristionchus pacificus* genomes using PROmer (version 3.07) [54] with default parameters. Macro-synteny was visualized using Circos (version 0.69) [55]. One linkage group (LG5) aligned almost exclusively to the *C. elegans* X chromosome. We genotyped 5 polymorphic markers from this linkage group in F1 hybrid males (from an APS4 x APS6 cross) confirmed it to be the X chromosome [20].

### Genome analyses

Orthologous proteins between *C. elegans* (PRJNA13758.WS264), *Haemonchus contortus* (HCON_v4, early access granted by Stephen Doyle), *P. pacificus* (El Paco v1), *Oscheius tipulae* (CEW1_nOt2) and *A. rhodensis* (chromosomal assembly) were identified through reciprocal best hit BLAST searches (BLASTp, “-evalue 0.01 -max_target_seqs 100, -outfmt 6”). Localisations of orthologous proteins were visualised using Circos plots (version 0.69) [55]. For *O. tipulae*, we used the correspondence of the genomic scaffolds to chromosomes [56]. For the visualisations of *O. tipulae* chromosomes, the scaffolds were collated in numerical order (the true order is currently not known). Gene (protein-coding and functional RNA) density was plotted for each chromosome using the R package karyoploteR (version 1.5.1)[57]. Protein-coding genes conserved between *A. rhodensis* and *Drosophila melanogaster* (GCF_000001215.4_Release_6_plus_ISO1_MT) were identified by performing reciprocal BLASTp searches (BLAST+, -evalue 0.01 -max_target_seqs 100, -outfmt 6). The localization of the conserved genes along the chromosomes was visualized using karyoploteR.

Variants (single nucleotide polymorphisms (SNPs) and insertions/deletions (InDels)) were identified using the three paired-end libraries for APS4 and the paired-end library for APS6, filtered as described above. Cleaned reads were aligned to the chromosomal assembly using bwa (version 0.7.12-r1039) [58] and the resulting SAM alignments were converted to BAM format and sorted by coordinate using Picard (version 2.14) SortSam, deduplicated using picard MarkDuplicates and the BAM files indexed using picard BuildBamIndex. The three APS4 libraries were merged prior to deduplication and indexing. Joint variant calling was performed using Samtools mpileup (version 1.4) [59] and the raw BCF output was filtered using bcftools view (version 1.4-16-g4dc4cd8) [60] and vcftools vcf-annotate (version 0.1.14, “-f +/d= 5/D= 10000/q= 20/Q= 15/w= 20/W= 30/c= 3,10/a= 2/1= 0.0001/2= 0/3= 0/4= 0.0001”) [61]. Intra-strain variants were defined as heterozygous polymorphisms occurring within one strain regardless of polymorphism at the same locus in the other strain. Inter-strain polymorphisms were defined as different genotypes between the two strains at the same locus. Intra- and inter-strain variant density was plotted along each chromosome using KaryoploteR [57]. The gene and variant densities of unanchored scaffolds (i.e. those not mapped to a linkage group) were not examined.

Gene ontology (GO) enrichment analyses were performed to examine possible GO terms found over- or under-represented in the X chromosome gene set versus the autosomal one. For each enrichment analysis, we used a two-tailed Fisher’s exact test (FDR < 0.05) implemented in the program Blast2GO (version 4.1.9) [42]. The list of GO terms found enriched or depleted in the test set was then reduced to the most specific terms.

### RNA extractions and transcriptomic analyses

#### L2 females, converted females and hermaphrodites

Female- and hermaphrodite-fated L2 larvae were isolated by using synchronized progeny populations generated by hermaphrodite mothers. Briefly, dauers (fated to become hermaphrodites) were isolated and allowed to develop into adult hermaphrodites. After ~12 h of egg laying the mothers were removed and the early eggs laid were left to hatch and grow until the L2 stage. During the L2 stage, females and hermaphrodites are distinguishable by their size and coloring: hermaphrodites are smaller, develop slower, are thinner and darker than females. Additionally, the female and hermaphrodite gonads are different in size during the mid-L1 stage, the female gonad being larger than that of the hermaphrodite [15, 18]. To convert hermaphrodite fated larvae into females, mid-L1 larvae with smaller gonads were isolated and grown with OP50-1 containing 200 nM dafachronic acid (DA) on NGM in individual wells of a 12 well culture plate. Once the larvae reached the L2 stage, the ones that had a female morphology were collected and used for RNA extraction. About 200 L2s of each sex were picked and transferred to an Eppendorf tube containing 200 µL of M9 buffer [22]. The nematodes were washed 2-3 times in M9 buffer, allowing them to sink to the bottom of the tube under gravity between each wash. After the final wash, the maximum amount of M9 was removed and 200 µL of Trizol was added to the tube. The tube was placed at −80 °C immediately. For RNA extraction, nematodes were first freeze-cracked in liquid nitrogen (2-3 times). Trizol was added to make up the volume to 500 µL and nematodes were shaken with a few sterile 0.5 mm glass beads on a BeadBeater homogenizer (20 s, 3 times with 30 s intervals). Subsequently, RNA was extracted using a standard chloroform approach and the pellet was dissolved in DEPC treated water and stored at −80 °C until further use.

#### Males and mixed stages samples

The same protocol was used to extract RNA from adult males and animals from various stages (mixed stages). To obtain RNA from males, about 500 young adult males were picked in a 1.5 mL centrifuge tube containing 0.5 mL of M9 and washed twice with the buffer. M9 was then replaced with 0.5 mL Trizol and the tube frozen at −80 °C. For the mixed staged nematodes, M9 was gently added to 5 culture plates (6 cm) containing a healthy population of nematodes. We avoided disturbing the bacterial lawn and naturally let the nematodes start to swim in the buffer. This method reduced bacterial contamination in the samples during harvesting. These nematodes were then collected into a centrifuge tube washed twice with M9, and then frozen using liquid nitrogen. Tissues were homogenized for 1 min using a probe homogenizer. After chloroform extraction, the RNA was dissolved in DEPC treated water and stored at −80 °C.

We generated three biological replicates for each RNA-seq condition (L2 females, L2 converted females, L2 hermaphrodites, males and mixed stages). RNA-seq was performed on the Illumina HiSeq2500 platform, generating a mean of 19.7 million 100 base read pairs per replicate. General assessment of the RNA-seq libraries was performed using FASTQC [62]. The raw reads from each library were preprocessed using Trimmomatic (version 0.36, “HEADCROP:15 SLIDINGWINDOW:5:20 MINLEN:20”) [63]. The RNA-seq aligner STAR (version 2.4.2, “–sjdbOverhang 84”) [64] was used to align the processed reads of each library to the primary scaffolded genome assembly. Transcript abundances were obtained using FeatureCounts (from the SubRead Package, version 1.5.0-p2) [65]. Differential expression between the L2 females, L2 converted females and L2 hermaphrodites (three comparisons) was assessed using the R package DEseq2 (version 1.18.1, [66]) following the standard procedure and generating diagnostic plots (as described in the DESeq2 documentation at https://bioconductor.org/packages/release/bioc/vignettes/DESeq2/inst/doc/DESeq2.html). An adjusted P-value of 0.01 and an absolute log2 fold change (FC) of 2 were used to define differentially expressed (DE) genes. Fold change of DE genes were plotted along the chromosomes using the package karyoploteR after lifting over annotations. Gene ontology (GO) enrichment analyses were conducted on the “down-regulated” and “up-regulated” genes separately for each comparison using the procedure explained above. Homologues of known sex determination genes were identified through reciprocal best hit BLASTp searches (BLAST+, “-evalue 0.01 -max_target_seqs 100, -outfmt 6”) using the *A. rhodensis* and *C. elegans* proteomes and analysed manually.

Global protein-coding gene expression of L2 females, L2 hermaphrodites, adult males and mixed stages was examined separately for each chromosome. The transcript abundances obtained using FeatureCounts (from the SubRead Package, version 1.5.0-p2) [65] were corrected by library size. The log2 of the global gene expression of each chromosome was plotted using ggplot2 in R. To determine if the genes of the X chromosome were significantly less expressed than those on the autosomes, we randomly sampled the same number of autosomal genes and X genes (600) and compared the sets using a Kruskal-Wallis test followed by Wilcoxon-Mann-Whitney tests between autosome-X pairs. The gene expression across a same chromosome in different replicates of a same condition was confirmed to be similar by performing Kruskal-Wallis tests.

### RNAi of the DM domain gene Arh.g5747

The DM domain gene Arh.g5747 was significantly up-regulated in hermaphrodites compared to females in both female and converted female samples at L2. To further investigate this sex-specific expression of Arh.g5747 we targeted the gene for down-regulation using RNA interference (RNAi). Target specific dsRNA was produced using a cDNA template. PCR amplification was performed using the following primers (Forward primer: 5’-*TAATACGACTCACTATAGGG*TCATCAACGAGCAGAGCCGAGA-3’, reverse primer: 5’-*TAATACGACTCACTATAGGG*TCCGCCTTCAGTGTTGGAGCT-3’) to amplify an 858 bp fragment of the transcript of Arh.g5747. The T7 promoter (shown in italics above) was included at the 5’ end of each primer to allow *in vitro* dsRNA synthesis. RNA extraction and cDNA synthesis were performed on ~300 adult hermaphrodite individuals as detailed above, with the exception that samples were subjected to repeated cycles of freeze-thawing instead of bead-beating. RNA was treated with DNase I (Sigma) to remove residual genomic DNA. cDNA synthesis was performed with 0.5 µg of RNA using random primers (Promega) and the MMLV reverse transcriptase enzyme (Promega) following the manufacturer’s instructions. The Arh.g5747 cDNA was then PCR amplified using GoTaq Green Master Mix (Promega), using approximately 200 ng of cDNA and 250 nM of each primer. PCR conditions were: an initial denaturation at 94 °C of 7 min, followed by 30 cycles of 94 °C for 15 s, 55 °C for 30 s and 72 °C for 60 s and a final extension of 10 min at 72 °C. After verification of the product size by gel electrophoresis, the amplicon was cleaned using the QIAquick PCR Purification kit (Qiagen), according to the manufacturer’s protocol and eluted in a final volume of 25 µL. dsRNA was *in vitro* transcribed by incubating approximately 200 ng of the cDNA template with 2 µl (40 U) of T7 polymerase (Promega), 20 µl of 5x T7 polymerase buffer (Promega), 10 µl of DDT (Promega),2.5 µl (100 U) of rRNAsin (Promega), 20 µl of 2.5 mM rNTPs (Thermo Fisher Scientific) and RNase free water to a final volume of 50 µl, for 4 h at 37 °C. The dsRNA product was size verified by gel electrophoresis and cleaned using the RNA clean-up protocol in the RNeasy^®^ Mini Kit (Qiagen). A mixture of short interfering RNAs (siRNA) was produced by digesting 5 µg of the Arh.g5747 dsRNA with ShortCut RNase III (NEB) for 20 min at 37 °C, cleaned by glycogen/ethanol precipitation and eluted in 20 µl of RNase free water, according to the manufacturer’s instructions. The RNAi mixture was produced by combining 5 µl of Arh.g5747 siRNA (approximately 100 ng), 4 ul of M9 buffer [22] and 1 µl (10% v/v) Lipofectamine^®^RNAiMax reagent (Invitrogen) and incubating at 25 °C for 20 min. Previously, we have shown that inclusion of the transfection reagent Lipofectamine dramatically improves RNAi efficiency in *A. rhodensis* [16]. For control injections Arh.g5747 siRNA was omitted from the mixture and replaced with additional M9 buffer.

For RNAi, young hermaphrodites (day 1 of adulthood) were immobilised on dried 2% agarose (w/v) pads in a small drop of undiluted halocarbon oil 700 (Sigma). The required injection mixture was loaded into pre-pulled microcapillary needles (Tritech Research) and microinjected into the gonad arms using an IM-300 Pneumatic Microinjector system with an oil hydraulic Micromanipulator (Narishige) using an injection pressure of 20 psi. Injected worms were rescued by adding a drop of M9 to the slide and moving them separately to a fresh 6 cm NGM plate seeded with *E. coli* OP50-1. The self-progeny from the injected hermaphrodites was sexed throughout the life of the mother. The mother was placed to a new plate every 24 h. Sex was determined according to the developmental rate, coloration and morphology of the larvae. Females were larger and whiter than hermaphrodites (dark and thin) due to their faster development. Males displayed a characteristic blunt tail.

## Results

### Genome characteristics

The scaffolded genome assembly spans 60.6 Mb in 440 scaffolds longer than 1000 bp. This span is smaller than that of *C. elegans* (100.2 Mb) but similar to that of other rhabditine nematodes (*Heterorhabditis bacteriophora*, 76.8 Mb; *Oscheius tipulae*, 59.0 Mb; *Caenorhabditis sulstoni*, 65.1 Mb). We predicted 11,570 protein coding genes and 833 unique non-coding RNAs. Previously sequenced rhabditine nematodes have been predicted to have more protein coding genes (*C. elegans*, 20,082; *H. bacteriophora*, 15,701; *O. tipulae*, 14,650). We have not explored the difference in coding gene content in *A. rhodensis* in detail. The top BLAST hits of *A. rhodensis* proteins were more likely to be from parasitic Strongylomorpha species (*Ancylostoma ceylanicum*, *Haemonchus contortus* and *Necator americanus*) than from *C. elegans*. This is consistent with molecular phylogenies derived from ribosomal RNA and small numbers of protein coding loci [7]. However, it conflicts with analyses based on larger protein-coding gene datasets which link *A. rhodensis* and the free-living *O. tipulae* to *Caenorhabditis* species. A majority of the proteins (10,449, 90%) was assigned at least one type of annotation (InterProScan signature, GO term, BLAST hit) and 8181 were decorated with at least one GO term. The non-coding RNAs included all the expected major classes (Supplementary Table 1). We also assembled and annotated the circular, 13,907 base pair mitochondrial genome (Supplementary Figure 1).

### Genetic map

We generated 95 F2-derived progeny Advanced Intercross Lines (AILs) from crosses between two polymorphic inbred strains of *A. rhodensis*, APS4 and APS6. We identified 1,052 polymorphic RAD-seq markers that clustered in 7 linkage groups (Table 3), presumably corresponding to the seven chromosomes in *A. rhodensis* identified by DAPI staining [20]. We anchored the genomic scaffolds of *A. rhodensis* to the genetic map to complete the sequence of each linkage group. Of the 1,052 markers and 636 scaffolds (> 200bp), 1,038 markers (~94%) and 143 scaffolds (~22%) were used to build the chromosomal assembly. The excluded scaffolds were generally short, and either lacked a RAD marker or had a marker that was not able to be placed. The anchored scaffolds represent 95.3% of the span of the scaffold span and contain 93.8% of the predicted proteins (Table 3).

**Table 2.** Genome assembly metrics of *Auanema rhodensis* and *Caenorhabditis elegans*.

| **Metric** | ***A. rhodensis***<br>**(scaffolds)** | ***A. rhodensis***<br>**(chromosomal**<br>**assembly)** | ***C. elegans***<br>**(PRJNA13758)\*** | ***O. tipulae***<br>**(CEW1_nOt2**<br>**)** |
| --- | --- | --- | --- | --- |
| **Number of scaffolds or chromosomes** | 636 | 6 autosomes<br>1 X<br>493 unplaced<br>scaffolds | 5 autosomes<br>1 X | 191 |
| **Assembly size or draft genome size**<br>**(Mb)** | 60.6 | 57.8 | 100.2 | 59.4 |
| **Number of scaffolds (> 200 bp)** | 636 | 7 | 6 | 191 |
| **Number of scaffolds (> 1,000 bp)** | 440 | 7 | 6 | 157 |
| **N50 (scaffolds > 1,000 bp) (bp)** | 556,081 | 8,804,062 | 17,493,829 | 1,203,411 |
| **Longest scaffold / chromosome (bp)** | 3,360,731 | 9,627,060 | 20,924,180 | 4,597,891 |
| **GC content (%)** | 32.2 | 32.2 | 35.44 | 44.53 |
| **Span of runs of Ns (>= 10 Ns) (bp)** | 915,180 | 928,780 | NA | 16310 |
| **Protein coding gene annotations** |  |  |  |  |
| **Number of genes (protein-coding)** | 11,570 | 10,861 | 23,629 | 14,938 |
| **Exons** |  |  |  |  |
| **Number of coding exons** | 135,144 | 130,644 | 189,079 | 127,820 |
| **Combined length of exons** | 16,655,465 | 15,869,469 | 39,400,137 | 20,438,569 |
| **Exon mean length (SD) (bp)** | 123.2 (131.1) | 121.5 (121.8) | 208.4 (263) | 159.9<br>(138.09) |
| **Exon median length (bp)** | 109 | 109 | 146 | 132 |
| **Minimum/maximum exon length (bp)** | 3 / 11,659 | 3 / 11,659 | 1 / 14,975 | 3 / 10,071 |
| **Introns** |  |  |  |  |
| **Number of of introns** | 123,693 | 119,812 | 200,020 | 112,925 |
| **Combined length of introns (bp)** | 16,862,509 | 16,308,478 | 79,153,867 | 17,941,595 |
| **Intron mean length (SD) (bp)** | 136.3 (521.3) | 136.1 (521.4) | 395.7 (962.3) | 158.9 (366.8) |
| **Intron median length (bp)** | 47 | 47 | 82 | 41 |
| **Minimum/maximum intron length (bp)** | 7 / 24,877 | 7 / 24,877 | 1 / 100,912 | 7 / 11,345 |

\* Metrics for *C. elegans* were calculated from the WormBase annotations (WormBase web site, http://www.wormbase.org, release WS262, July 2018).

**Table 3.** Characteristics of the *A. rhodensis* genetic map.

| **Linkage group** | **Number of RAD-seq markers** | **Homozygous APS4/APS4 frequency** | **Heterozygous APS4/APS6 frequency** | **Homozygous APS6/APS6 frequency** | **Assembly length (bp) \*** | **Number of protein-coding genes** |
| --- | --- | --- | --- | --- | --- | --- |
| **LG1** | 109 | 0.30 | 0.35 | 0.35 | 8,489,927 | 1,538 |
| **LG2** | 149 | 0.29 | 0.37 | 0.34 | 9,627,060 | 1,760 |
| **LG3** | 184 | 0.21 | 0.38 | 0.51 | 8,741,542 | 1,748 |
| **LG4** | 143 | 0.35 | 0.39 | 0.36 | 8,804,062 | 1,586 |
| **LG5X** | 92 | 0.04 | 0.88 | 0.08 | 3,488,253 | 604 |
| **LG6** | 185 | 0.35 | 0.39 | 0.26 | 9,421,540 | 1,871 |
| **LG7** | 190 | 0.23 | 0.42 | 0.35 | 9,306,279 | 1,754 |
| Overall | 1052 | 0.26 | 0.42 | 0.32 | 57,878,663 | 10,861** |

\* Length in chromosomal assembly after anchoring the scaffolds onto the genetic map.
\*\* 93% of total number of protein coding genes.

### Macrosynteny with *C. elegans* and identification of the X chromosome

*A. rhodensis* chromosomes are smaller than those of *C. elegans* (mean 8.3 Mb for *A. rhodensis* compared to 20.1 Mb for *C. elegans*), and one chromosome, LG5, is less than half the average size. To explore the origins of the changed complement of chromosomes and reduced size, we aligned the *A. rhodensis* protein-coding gene set to the *C. elegans* one (Figure 2). While there was a minor background of between-chromosome translocation, most chromosomes had congruent gene sets. The majority of loci on three *A. rhodensis* linkage groups (LG1, LG6 and LG7) mapped to single *C. elegans* chromosomes (V, IV and II, respectively), suggesting one-toone correspondence. For *A. rhodensis* LG2, LG3 and LG4 we observed mapping to two or more chromosomes. Thus, LG2 of *A. rhodensis* combines components on *C. elegans* chromosomes III and X, LG3 combines elements of I and III, and LG4 combines parts of I, III and X. LG5, the smallest *A. rhodensis* chromosome, mapped almost entirely to the X chromosome of *C. elegans*, but components of *C. elegans* X were also found on LG2 and LG4 (Figure 2).

We confirmed that *A. rhodensis* LG5 was the X chromosome by genotyping F1 hybrid APS4/APS6 males at polymorphic loci spread across the linkage groups [20]. Hybrid males were heterozygous for all inter-strain polymorphic loci with the exception of those loci located in LG5, which were hemizygous in males [20]. Since males produce only one kind of functional sperm, with the X chromosome [17], all F1 sons carried the paternal genotype in the X chromosome [20].

### Chromosomal rearrangements

To further understand chromosomal rearrangements which took place in the branch leading to *A. rhodensis*, we analysed the synteny relationships of loci conserved between *A. rhodensis* linkage groups and the chromosomal assemblies of *C. elegans*, *H. contortus, O. tipulae* and *Pristionchus pacificus* (Figures 1 and 3, Supplementary Figure 2).

**Figure 1.**
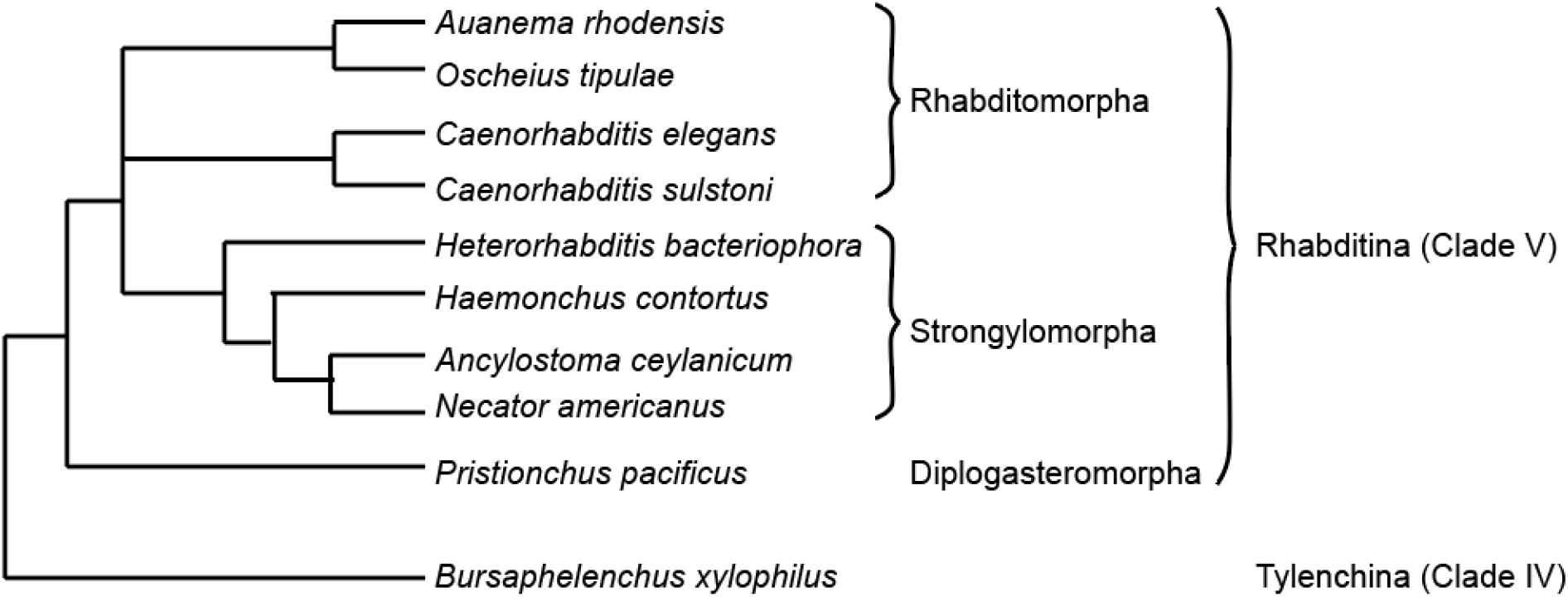
The relationship of *Auanema rhodensis* to other rhabditid nematodes. The phylogeny of the nematode species discussed in this analysis. *A. rhodensis* and *O. tipulae* are sister taxa in analyses based on multiple protein coding gene and ribosomal RNA loci [7, 46,67], but their relationships to *Caenorhabditis* and the parasitic Strongylomorpha differ depending on the datasets analysed.

**Figure 2.**
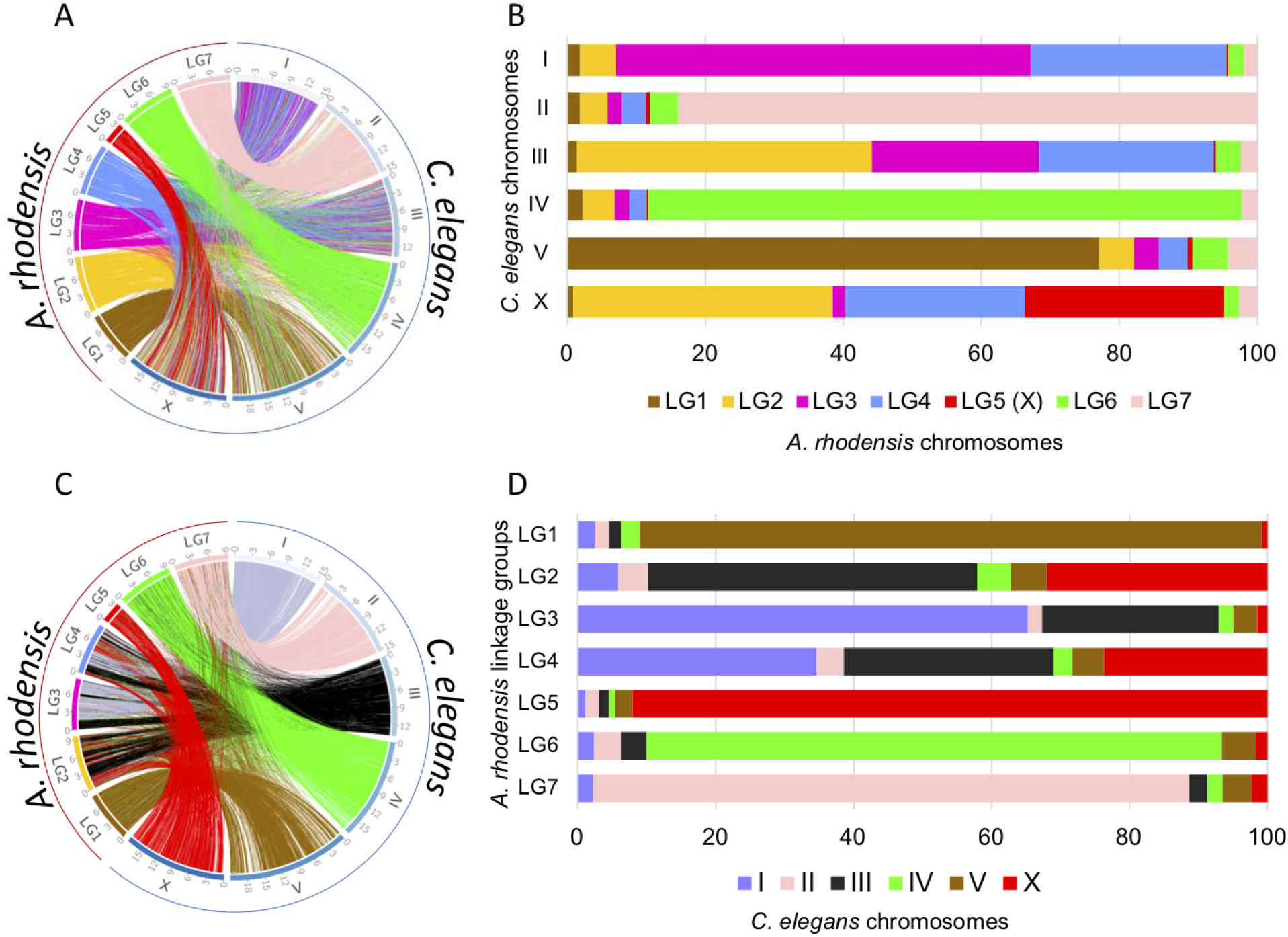
Synteny relationships between chromosomes of *Auanema rhodensis* and *Caenorhabditis elegans*. Location (A and C) and proportion (B and D) of orthologous protein-coding genes between *C. elegans* and *A. rhodensis* coloured according to *A. rhodensis* (A and B) or *C. elegans* (C and D) chromosomes.

**Figure 3.**
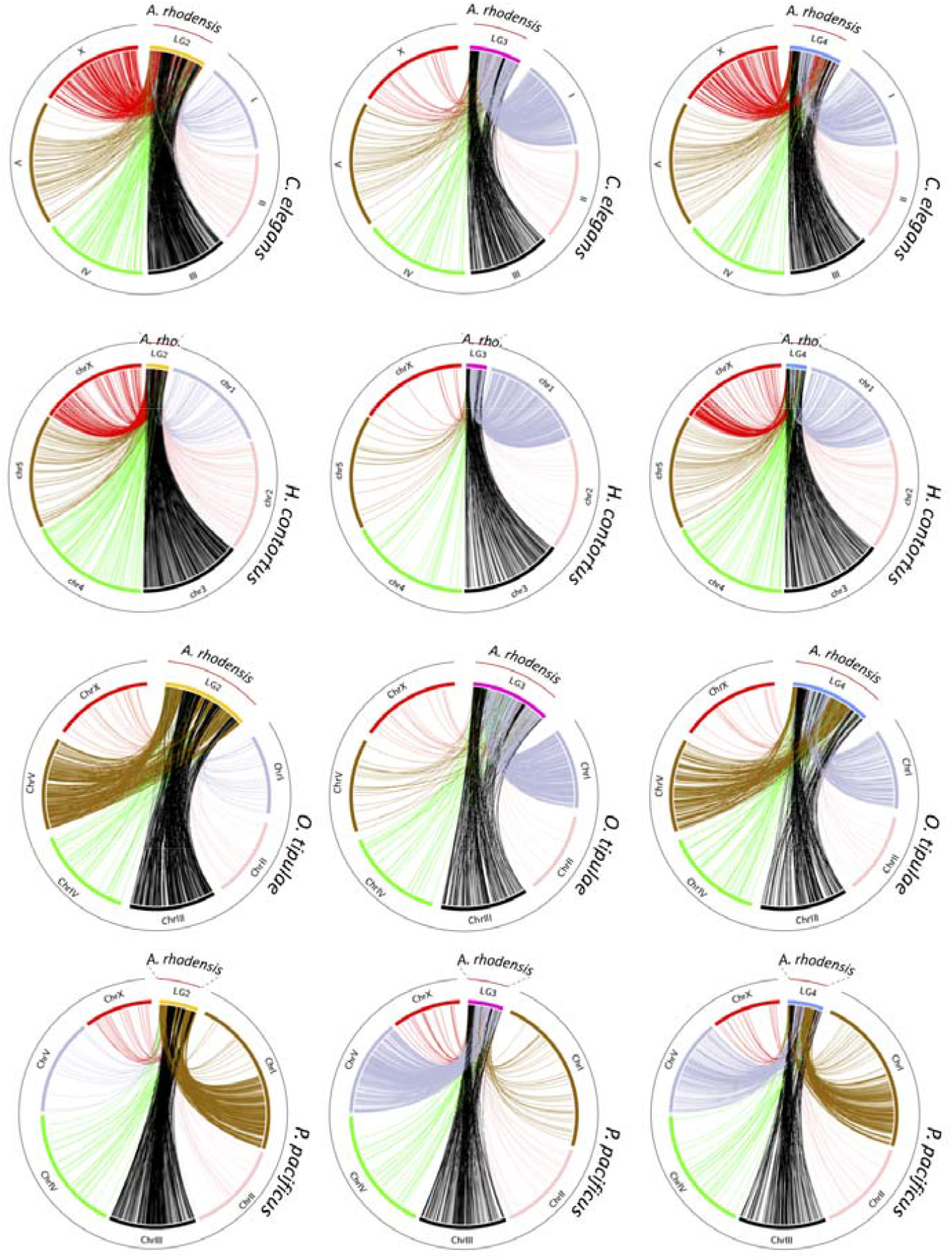
Macrosynteny relationships of *Auanema rhodensis* linkage groups LG2, LG3 and LG4. The circos plots show macrosynteny relationships of (columns) LG2 (yellow), LG3 (pink) and LG4 (blue) of *A. rhodensis* to (rows) the chromosomal genomes of C. elegans, *H. contortus, O. tipulae* and *P. pacificus*, based on the mapping of presumed orthologues between the species.

*A. rhodensis* LG1 (Ar_LG1) contained loci that had orthologues on a single chromosome in *C. elegans* and *H. contortus* (chromosomes V/5; Ce_V, Hc_5) and on the left arm of *P. pacificus* chromosome I (Pp_IL) (Figure 1 and Supplementary Figure 2). Comparing to *O. tipulae*, the orthologues of loci on Ar_LG1 were on chromosome X (Oti_X) (Supplementary Figure 2). *A. rhodensis* LG6 and LG7 had similar single-chromosome counterparts in the other species analysed. Orthologues of loci on Ar_LG6 were found on Ce_IV, Ot_IV, Hc_4 and Pp_IV. Orthologues of loci on Ar_LG7 were found on Ce_II, Ot_II, Hc_2 and Pp_I (Figure 1 and Supplementary Figure 2).

Orthologues of loci on Ar_LG2 were found on two distinct chromosomes in *C. elegans*, *O. tipulae* and *H. contortus* (Ce_III, Hc_3, Ot_III and Ce_X, Hc_X, Ot_V). On Ar_LG2, these loci were partially segregated into blocks with different chromosomal locations in the other species (Figure 3, first three rows). In *P. pacificus*, these same blocks of loci had orthologues segregated on the right arm of chromosome 1 (Pp_IR) and on Pp_III. We concluded that Ar_LG2 was the product of fusion and rearrangement of a fragment or fragments of an ancestral chromosome represented by Pp_IR, Ot_V, part of Ce_X and part of Hc_X and an ancestral chromosome now present as parts of Pp_III, Ot_III, Ce_III and Hc_3. The presence of interspersed, extended blocks of loci that appeared to derive from the same ancestral chromosome suggested that the rearrangement was relatively recent, as the processes of intrachromosomal inversion, known to be very rapid in rhabditine nematodes [68], had not yet mixed up these blocks of genes.

Analyses of Ar_LG3 and Ar_LG4 identified similar patterns of breakage and fusion. Ar_LG3 contained two sets of distinct blocks of loci, one that had orthologues on Ce_III, Ot_III, Hc_3, and Pp_III, and a second that had orthologues on Ce_I, Ot_I, Hc_1, and Pp_V (Figure 3, middle column). Ar_LG4 had three sets of blocks of loci with orthologues on three chromosomes in other species: one set on Ce_III, Hc_3, Ot_III and Pp_III, one on Ce_X, Hc_X, Ot_V and Pp_ IR, and one on Ce_I, Hc_I, Ot_I and Pp_V.

### The evolutionary history of the X chromosome

The *A. rhodensis* X chromosome (LG5X) was the smallest chromosome (3.6 Mb) and had the lowest number of protein-coding genes (604, 5.5% of the total). X chromosomes differed markedly between species, but each contained orthologues of loci found on Ar_LG5X. For example, when compared to the most distantly related species, *P. pacificus* (Figure 4), the majority (82%) of the homologues of genes on Ar_LG5X were found on Pp_X (Figure 4A, Supplementary Figure 2C). However, Pp_X was five times as big (16 Mb) and contained 2,998 protein coding genes (11.7% of all predicted *P. pacificus* genes).

**Figure 4.**
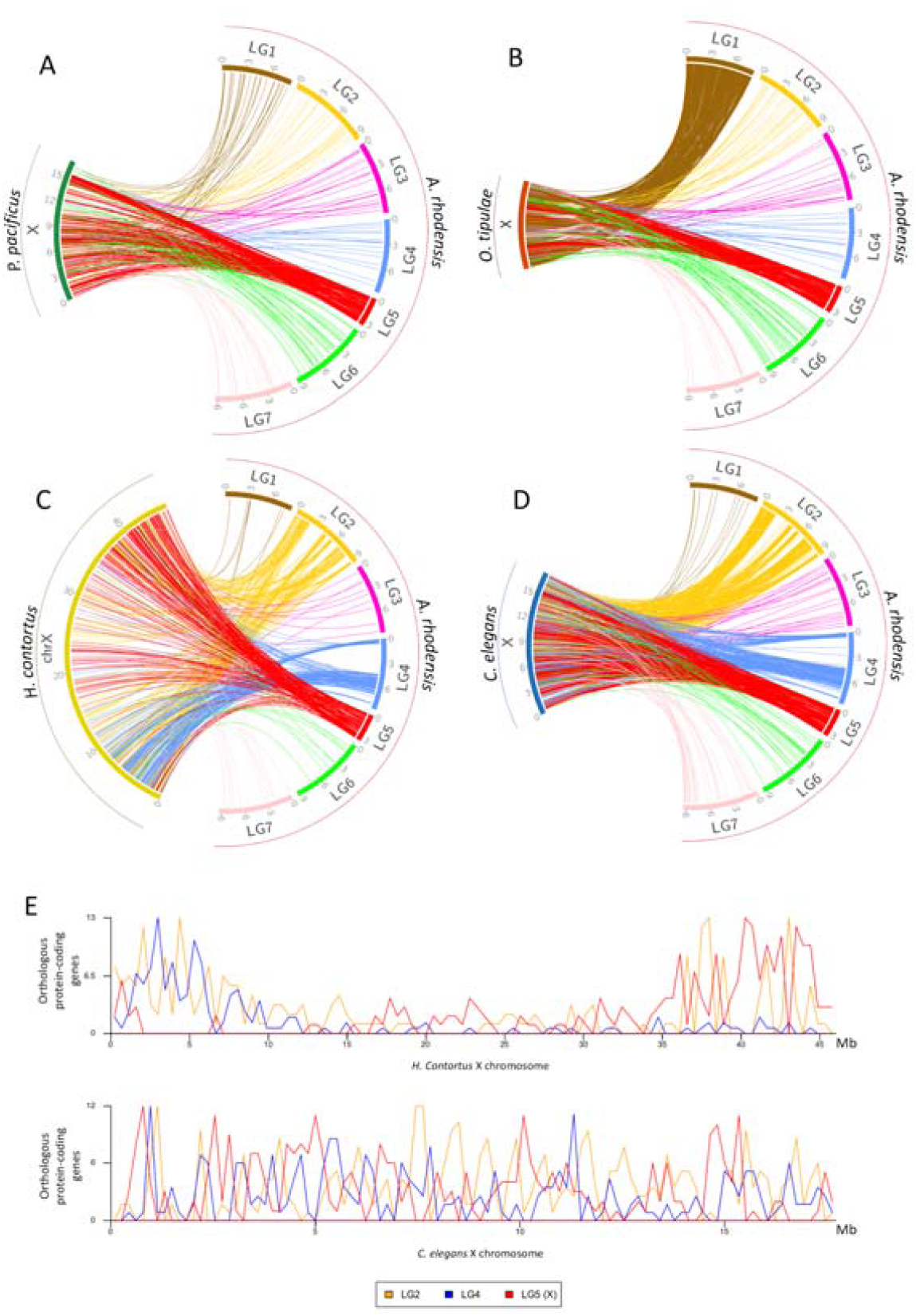
Macrosynteny relationships of rhabditine X chromosomes. For four rhabditine nematodes for which chromosomal genome assemblies or scaffold allocations to chromosomes (*O. tipulae*) are available, we mapped the location of their X-linked genes to the *A. rhodensis* genome (**A**: *Pristionchus pacificus*; **B**: *O. tipulae*; **C**: *Haemonchus contortus*; **D**: *Caenorhabditis elegans*). **E** Distribution of mappings to *A. rhodensis* chromosomes of X-linked genes in *H. contortus* (upper) and *C. elegans* (lower). The X of *O. tipulae* is represented as a concatenation of all the scaffolds belonging to the X chromosome; their order is arbitrary (scaffold number).

*C. elegans* and *H. contortus* X chromosomes shared a striking pattern of macrosynteny with *A. rhodensis*. Mapping homologues from the *C. elegans* X to *A. rhodensis*, there were three distinct blocks of synteny on Ar_LG4 (Figure 4D), and 5 blocks of synteny on Ar_LG2 (Figure 4D). These same blocks were observed in comparisons with *H. contortus* (Figure 4C). Intriguingly, while the *C. elegans* orthologues of Ar_LG5X loci were evenly distributed across Ce_X (Figure 4E), the mapping of Ar_LG5X orthologues on the *H. contortus* X was partitioned, albeit less clearly than the blocky mapping in *A. rhodensis*. Ar_LG4 matches were clustered on the left end of Hc_X and Ar_LG5X matches on the right (Figure 4E). Ar_LG2 matches were more evenly distributed, although we can note a clustering on both ends of Hc_X. As discussed above, Ar_LG2 and Ar_LG4 may have originated through chromosome breakage and rearrangement. The segregation of Ar_LG5X-like and Ar_LG4-like regions on the *H. contortus* X may reflect conservation of ancestral synteny that has not been homogenised by within-chromosome rearrangement. The contrast between Ce_X and Hc_X, two chromosomes that otherwise appear highly homologous, suggested that either intrachromosomal rearrangement was been much more active in the lineage leading to *C. elegans* or that *A. rhodensis* and *H. contortus* shared a more recent common ancestor. *A. rhodensis* orthologues of genes on the *O. tipulae* X chromosome were found on Ar_LG5X and Ar_LG1 (Figure 4B). This pattern was different from the one shared by mappings to *P. pacificus*, *H. contortus* and *C. elegans* and may reflect a novel trajectory of X chromosome evolution in the branch leading to *O. tipulae*.

### Contrasting patterns of genome structure between the X chromosome and the autosomes

We explored large-scale patterns of genome structure and evolution across the *A. rhodensis* genome. In *C. elegans*, conserved genes are more frequently found in the centres of chromosomes and are rarer in autosomal chromosome arms [69, 70]. However, in *A. rhodensis* the gene density and localisation of genes with orthologues in *Drosophila melanogaster* across chromosomes was uniform (Figure 5A). LG5X had a lower gene density than the autosomes (one protein-coding gene per 5.8 kb on LG5X compared to 5.3 kb ± 0.2 kb on the autosomes), and fewer conserved genes were present on LG5X (Figure 5A, Supplementary Table 2). Most strikingly, none of the nearly 500 tRNA loci were on LG5X (Figure 5B).

**Figure 5.**
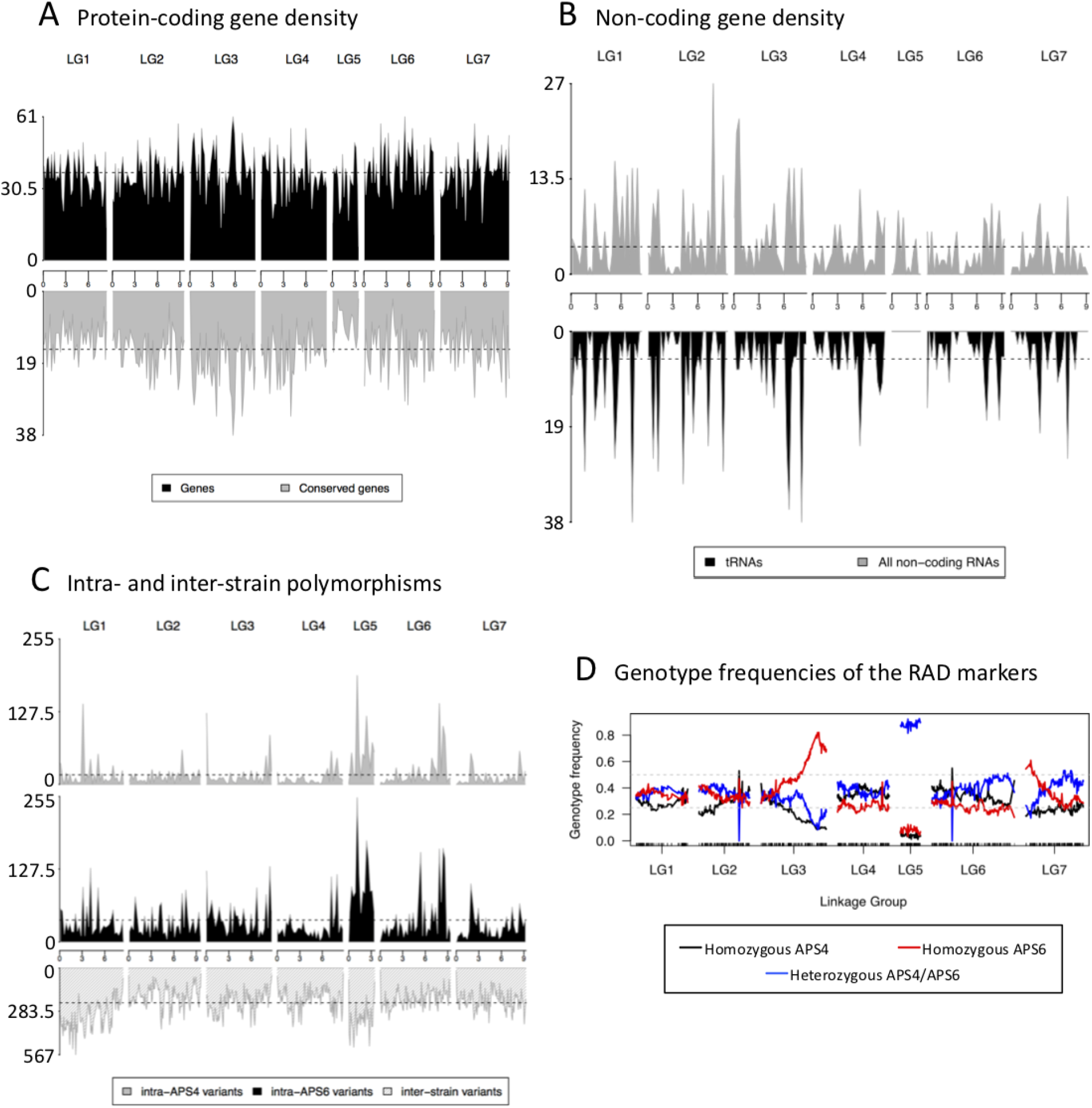
Contrasting genomic patterns between the X chromosome and the autosomes. (A) Distribution and conservation of protein-coding genes across *A. rhodensis* chromosomes. Density of *A. rhodensis* protein coding genes (upper panel) and conserved genes between *A. rhodensis* and *D. melanogaster* (lower panel) along each linkage group using a 200,000 bp window size. Overall, 4,544 conserved genes were identified between *A. rhodensis* and *D. melanogaster*. (B) Localisation of the annotated non-coding genes (upper panel) and transfer RNAs (tRNAs) (lower panel) along each linkage group using a 300,000 bp window size. No tRNAs were found on the X chromosome (LG5). (C) Patterns of variation across two inbred strains of *A. rhodensis*. Variant density along each chromosome in 250,000 base windows for the within-strain variants (upper panels) and 100,000 base windows for the between-strain variants (lower panel). (D) Genotype frequencies across *A. rhodensis* chromosomes. Black and red lines represent the frequencies of RAD sites homozygous for the APS4 or for the APS6 allele, respectively, in the 95 genotyped F2 AILs. Blue lines represent heterozygous genotypes. More than 80% of the progeny samples were heterozygous for the X chromosome (LG5X). The black ticks on the x-axis show the positions of the 1052 mapped RAD markers.

A gene ontology analysis comparing the X *versus* the autosomal gene sets found that the GO terms ‘translation’, ‘ribosome’, ‘nucleic acid binding’, ‘intracellular membrane-bounded organelle’ and ‘hydrolase activity’ were under-represented on LG5X compared to the autosomes (Supplementary Table 3). The process governing the ‘neuropeptide signaling pathway’ was found enrich on LG5X (Supplementary Table 3).

Although the strains used in this study were inbred, we expected to observe a low level of within-strain heterozygosity due to incomplete inbreeding. We found that strain APS6 had higher heterozygosity than APS4, probably because APS6 underwent less inbreeding than APS4 (11 rounds of bottlenecking *versus* 50) (Figure 5C, Supplementary Table 4). While the overall frequency of variants was different in the two strains, the distribution of these variants across the genome was similar (Figure 5C), including shared chromosomal regions with higher natural variability. The LG5X chromosome displayed more within-strain variation than the autosomes, probably due to the atypical inheritance of this chromosome.

The genetic map displayed deviation from expected Hardy-Weinberg equilibria in several regions. We found that almost all RAD markers for LG5X were heterozygous across all 95 samples (Figure 5D). This distorted pattern of X heterozygosity can be explained by the fact that the AIL RAD data were derived from F2 hermaphrodite progenitors left to propagate for 3-10 generations, and there is no X recombination in hermaphrodites [20]. We also observed a high frequency of homozygous markers for APS6 alleles at the right end of LG3 (Figure 5D), suggesting that APS6 alleles had been positively selected in the culture conditions tested or possibly that segregation distorters were present. Less extreme deviation from expected equilibrium was also observed at the left end of LG7. These deviations were not explored further.

The different dosage of X chromosomes in females and hermaphrodites compared to males results in a requirement for dosage compensation for X-linked genes. In turn, genes on the X often have distinct overall levels of expression compared to genes on autosomes. In *C. elegans* hermaphrodites the relative expression level of genes on the X varies through development from 1.29 fold higher than genes on autosomes in the L2 to 0.52 fold in adults [71]. This change is likely to be associated with the exclusion of genes with germline expression from the *C. elegans* X chromosome and the growth of the germline in adults [72, 73]. We examined global protein-coding gene expression in L2 females, L2 hermaphrodites, adult males and mixed stages and compared autosomal and X-linked genes (Figure 6). After correction for library size, gene expression from each autosome was found to be similar both between autosomes and across lifecycle stages. However, and unlike *C. elegans*, genes on *A. rhodensis* LG5X showed consistently lower expression than those on the autosomes, even in the L2 stage (Wilcoxon Mann-Whitney, p-value <= 1.0e-11 in all conditions and replicates).

**Figure 6.**
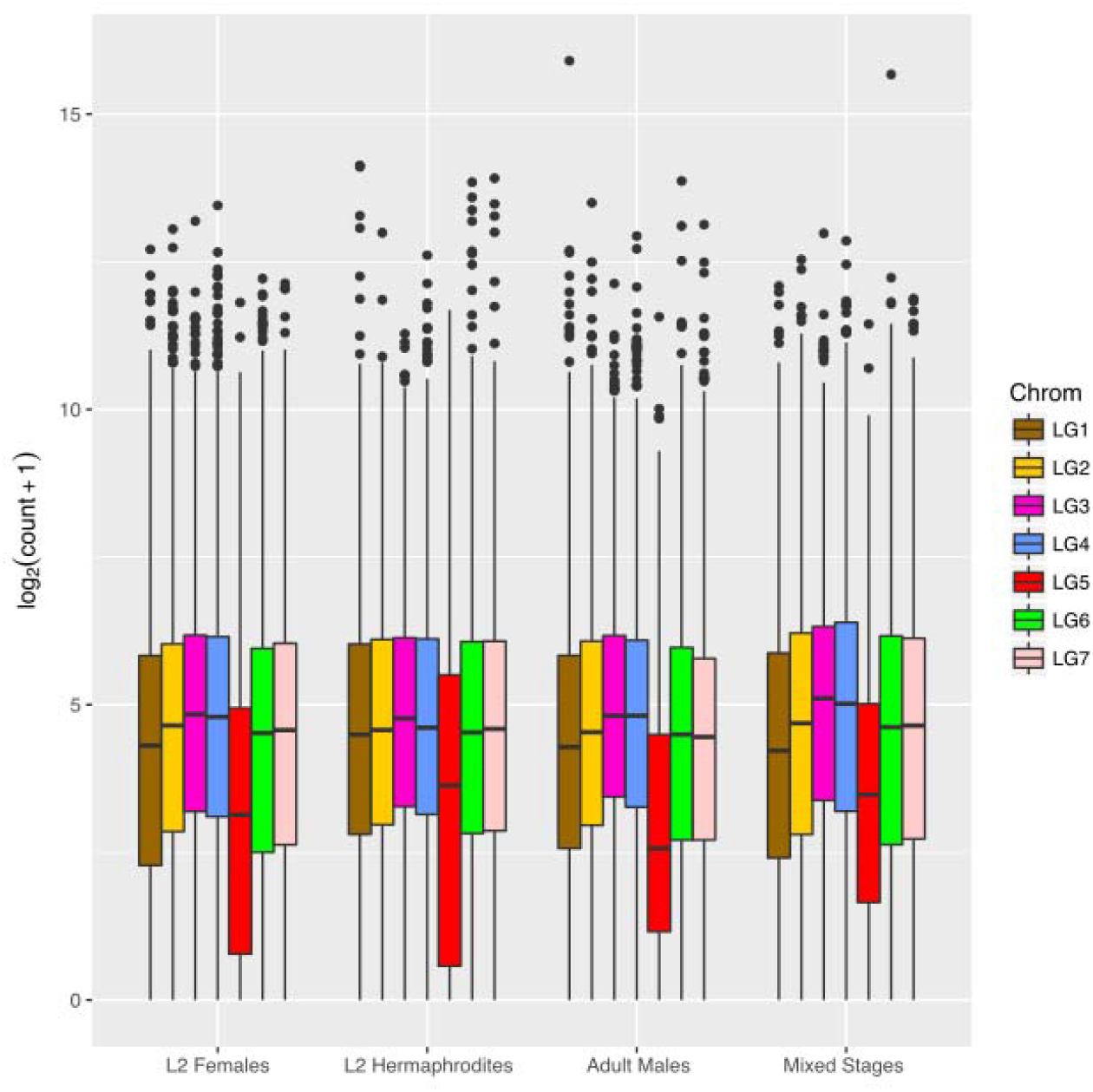
Expression of genes on the *A. rhodensis* X chromosome is generally lower than those on autosomes. Boxplots of the log2 normalized expression of the genes located on each linkage group of *A. rhodensis* in different sexes and stages in single replicate libraries. The expression levels were normalized by library size and log2-transformed. LG5 (in red) is the X chromosome. Boxplots for all libraries are represented in Supplemental Figure 3. This plot was generated using the R package ggplot2 [74].

### Transcriptomic identification of loci associated with sexual morph development

We compared gene expression in developing XX L2 larvae to identify loci that may be associated with the different sexual morphs of *A. rhodensis*. We generated replicate RNA-seq datasets from L2 fated to become hermaphrodites, L2 fated to become females, and L2 from hermaphrodite-fated nematodes that were converted to females by treatment with DA (converted females). Using standard thresholds (absolute log2(Fold Change) >= 2, FDR <0.01), we found 2,422 (21%) of the predicted genes were differentially expressed (DE) between L2 hermaphrodites and L2 females. Slightly fewer genes (2,121,18%) were DE between L2 hermaphrodites and L2 converted females. Most of the genes found to be DE between females and hermaphrodites and between converted females and hermaphrodites were the same (Figure 7A). The genes more expressed in females and converted females compared to hermaphrodites were enriched in GO terms related to translation, protein synthesis, ribosomal function, gonad and embryo development and structural constituents of the cuticle (see Supplementary Data 1 for the list of complete terms). Genes more expressed in hermaphrodites were enriched in few GO terms, with only “structural constituent of cuticle” in common between both comparisons. DE genes were distributed across the *A. rhodensis* genome, with no enrichment or depletion on LG5X (Figure 7B).

**Figure 7.**
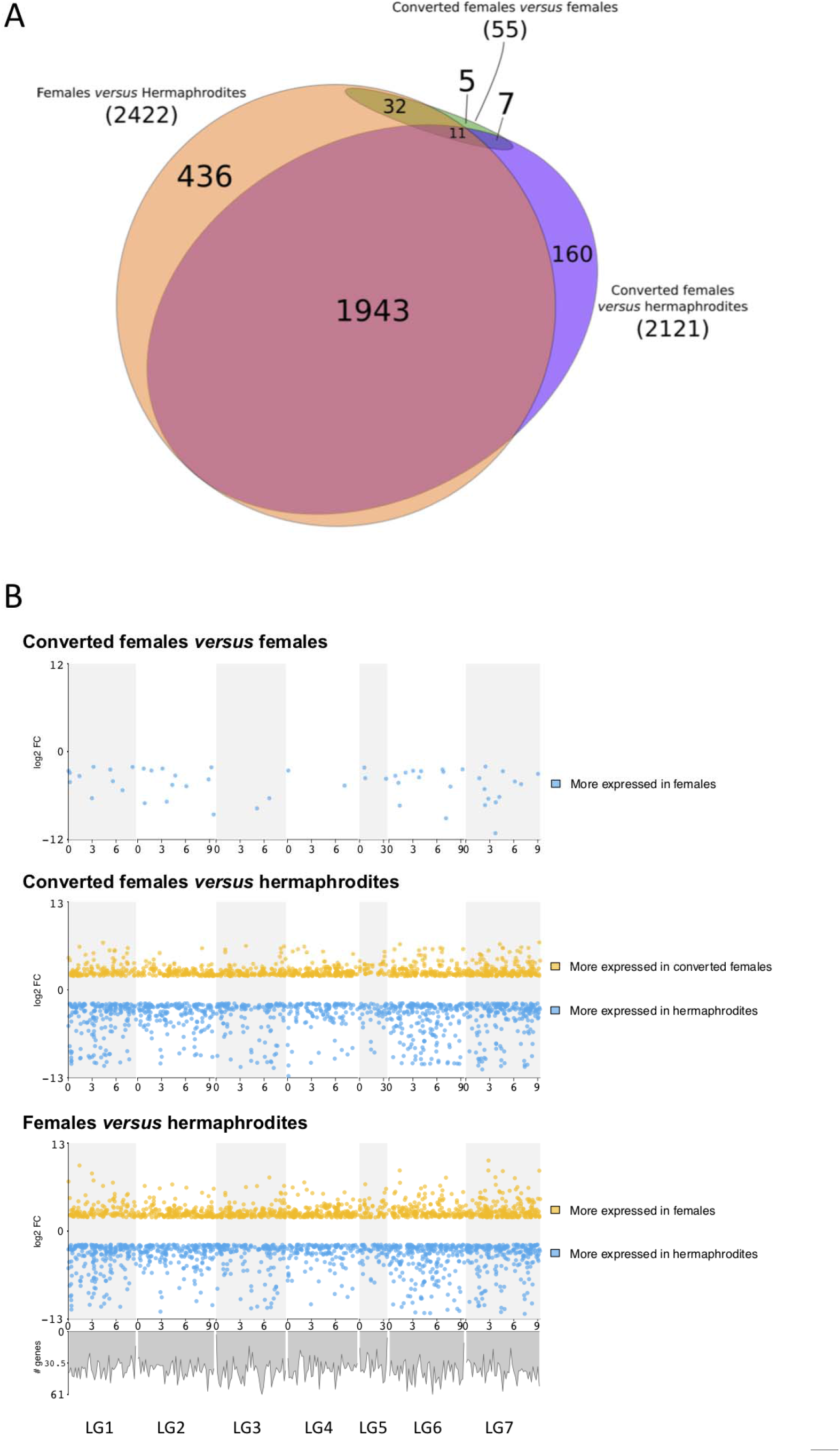
Analysis of differential gene expression in *Auanema rhodensis* XX nematodes. A. Differentially expressed genes in *A. rhodensis* XX L2 larvae. Most genes found to be DE between female L2 and hermaphrodite L2 were also differentially expressed between the converted female L2 and the hermaphrodite L2. Few DE genes were found between the female L2 and converted female L2. Of these, 32 had similar expression in hermaphrodite L2. B Distribution of DE genes (absolute log2(FC) >= 2, FDR <0.01) along the chromosomes of *A. rhodensis*. Upper panel: Female L2 *versus* converted female L2; middle panel: hermaphrodite L2 *versus* converted female L2; lower panel: female L2 *versus* hermaphrodite L2. A plot of the density of all genes along the genome (see Figure 5C) is shown under the lower panel.

Some *A. rhodensis* orthologues of *C. elegans* sex determination genes were DE between female (normal or converted) and hermaphrodite L2s. The known sex determination genes Arh.g5696-*gld-1* and Arh.g4999-*tra-1* were 200 and 4 times more expressed in females, respectively. The precise roles of *tra-1* and *gld-1* in the sex determination in *A. rhodensis* are not yet known. A number of *daf* (dauer formation) genes (Arh.g6122-*daf-11*, Arh.g7695-*daf-16*, Arh.g7696-*daf*-like) were expressed at higher levels in hermaphrodite L2 compared to female or converted female L2, consistent with the obligate transition through dauer of hermaphrodite-fated nematodes.

The DM (*doublesex*/*mab-3*) domain transcription factor Arh.g5747 (*dmd-10*/*11*-like) was found to be more than 200 times more expressed in hermaphrodite L2 than in female or converted female L2. To investigate the role of this locus in the decision between female and hermaphrodite sexual fate in *A. rhodensis*, we downregulated Arh.g5747 by injecting RNAi in young hermaphrodites (first day of adulthood). If Arh.g5747 is required for determining hermaphrodite fate, we would expect to see more female progeny from injected hermaphrodite mothers. Indeed, downregulation of Arh.g5747 in 8 hermaphrodite mothers resulted in more female progeny than control injections performed on 9 hermaphrodites (Wilcoxon Mann-Whitney test, W = 67, p-value = 0.001563, Table 4). Thus, this DM domain locus may drive *A. rhodensis* hermaphrodite fate, either by inhibiting a female induction signal or through positive upregulation of a hermaphrodite-inducing pathway.

**Table 4.**
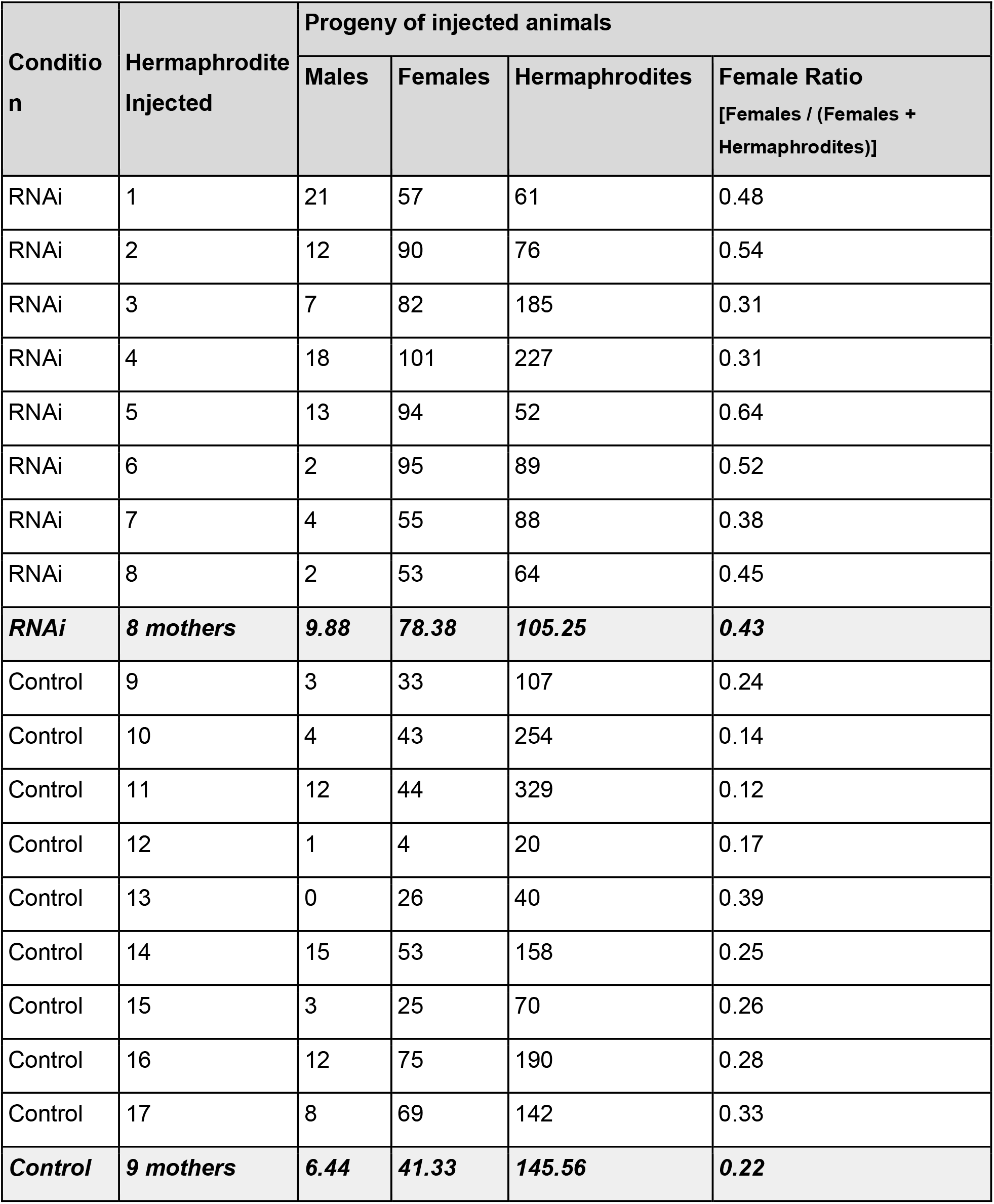
Inhibition of the DM transcription factor Arh-g5747 (dmd-10/11) by RNAi in hermaphrodite mothers results in more female progeny.

The comparison of the L2 females and the L2 converted females is particularly interesting for identifying genes or mechanisms involved in the hermaphrodite-female decision. Gene expression in L2 females and L2 converted females was strikingly similar. Only 55 genes (0.5%) were found to be more highly expressed in females compared to converted females (Figure 7A and 7B). No genes were found to be significantly less expressed in females compared to converted females. This result is surprising, since the female-inducing treatment (DA) was applied at the L1 stage, when sexual fate has already been decided, and the transcriptome was sampled less than 24 h after DA application. Functional annotation of these DE genes revealed several whose *C. elegans* homologues are involved in embryogenesis and developmental processes. Three chondroitin proteoglycan genes (Arh.g2548, Arh.g5439, Arh.g2211) were more expressed in female L2, and were also DE between female L2 and hermaphrodite L2. Chondroitin proteoglycans are important for embryonic cell division and vulval morphogenesis. More strikingly, we identified the homologues of the zinc-finger genes *mex-1* (required for germ cell formation, and somatic cell differentiation in the early embryo in *C. elegans*) and *pos-1* (essential for proper fate specification of germ cells, intestine, pharynx, and hypodermis in *C. elegans*). Both these genes had very low expression in converted female L2 and hermaphrodite L2. As these genes are maternally supplied in *C. elegans*, this supports a model where the decision between female and hermaphrodite fates is, at least partially, maternal.

## Discussion

*Auanema rhodensis* is a rare example of a three-sexed animal. Here we sequenced the *A. rhodensis* genome and used a linkage map to construct a chromosomal assembly. At 60 Mb, the *A. rhodensis* genome is smaller than that of *C. elegans* (100 Mb), but within the range (55-160 Mb) of other free-living rhabditomorph nematodes. We predict only 11,570 protein coding genes, many fewer than the 23,000 identified in *C. elegans*, and fewer than would be predicted from the reduction in genome size alone. Whether the reduced gene count is correct, or a reflection of the much more detailed analysis of the *C. elegans* genome, remains to be assessed.

It is known that mating system can influence genome size. In the *Caenorhabditis* clade, selfing (hermaphrodite) species have smaller genomes than their outcrossing sister species [75, 76], but these differences are in the order of 10%. However, other free-living and entomopathogenic rhabditomorphs have genomes smaller than *C. elegans*, and the related animal-parasitic Strongylomorpha have much larger genomes (250 – 700 Mb). Detailed understanding of the evolutionary drivers of genome size in this group awaits additional, dense sampling across the Rhabditomorpha.

While there is little shared gene order between nematode species, we identified strong macro-syntenic patterns between *A. rhodensis, P. pacificus, H. contortus, O. tipulae* and *C. elegans*. These patterns allow us to propose a preliminary model of the evolution of chromosomes in the Rhabditina, which includes Diplogasteromorpha (*P. pacificus*), Strongylomorpha (*H. contortus*), and Rhabditomorpha (*A. rhodensis, O. tipulae* and *C. elegans*). It has long been noted that the majority of rhabditine nematodes have a karyotype of n= 6, with an XX:XO sex chromosome system [77]. While there are deviations from this pattern, including the trioecious *A. rhodensis*, with n= 7, and the variously parthenogenetic *Diploscapter* species with n=1 to n=9, the phylogenetic perdurance of this karyotype is striking. Using loci identified as orthologues in each species pair, we could identify six putative ancestral macrosynteny groups (Figure 8) and also map the macrosyntenic changes that may have given rise to present day karyotypes.

**Figure 8.**
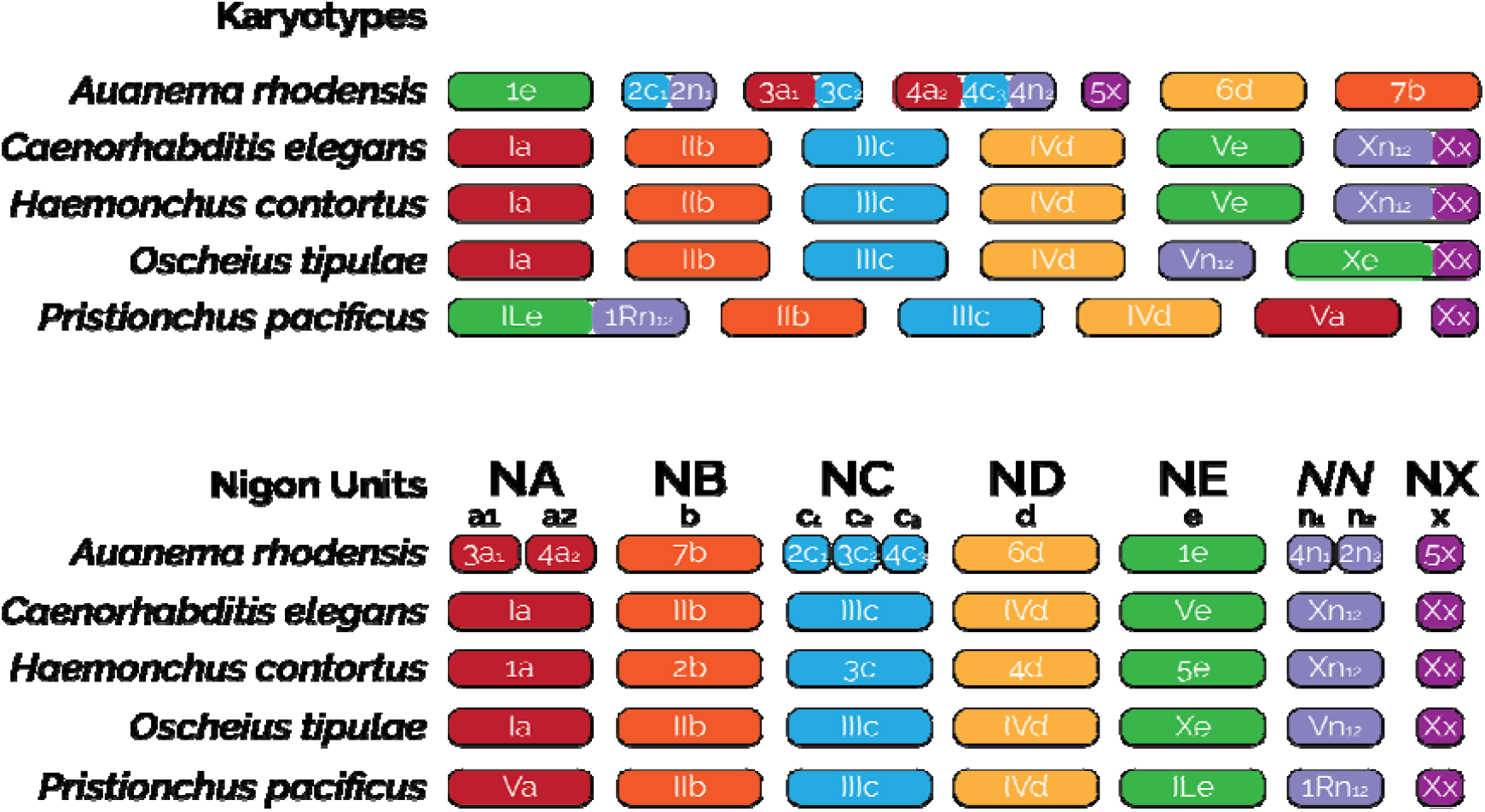
Nigon units and the evolution of rhabditine chromosomes. The different colors indicate orthologous chromosomes/chromosomal parts belonging to different Nigon units. Nigon unit ‘*NN*’ may be part of NX or NE or a separate unit as depicted here. Reshuffling within chromosomes is not depicted.

We call these ancestral linkage groups Nigon units, a name coined by Matt Rockman (personal communication) in analogy with the Muller units of *Drosophila* chromosomes. Some of these Nigon units have been transmitted intact through Rhabditina, while others have undergone rearrangement. Where a rearrangement has resulted in the fusion of (parts of) Nigon units, in most cases the dynamic processes of intrachromosomal rearrangement, which are very active in rhabditine nematodes [68], have acted to mix up the genes originally derived from different units. In other cases, either because the fusions were more recent or because the processes of intra-chromosomal rearrangement are less active, the sets of loci from distinct Nigon units are found as blocks in the fusion chromosome. We define six Nigon units, NA, NB, NC, ND, NE and NX, as well as an additional NN unit which we are currently unable to place (Figure 8). It could originally link to NE or NX, or be a unit of its own (which would imply seven Nigon units in total).

*A. rhodensis* LG1, LG6 and LG7 represent chromosomal units that have survived largely intact through rhabditine evolution. However, the *A. rhodensis* X chromosome (LG5X) was formed from a subset of the loci now found on the *C. elegans* X chromosome, and *A. rhodensis* LG2, LG3 and LG4 are the products of major interchromosomal rearrangement events. For example, NA is intact in *C. elegans* (Ce_I), *H. contortus* (Hc_1), *O. tipulae* (Ot_I) and *P. pacificus* (Pp_V), but underwent fission in the *A. rhodensis* lineage, with subsequent fusion forming two hybrid chromosomes. *A. rhodensis* LG3 is formed largely from loci from part of NA (subset a1) and NC (c2), while LG4 includes loci from NA (a2), NC (c1) and NN (n1). Overall, compared to the other four species with chromosomal assemblies (or chromosome-allocated scaffolds), The fission/fusion event(s) may be directly associated with the origin of the novel n= 7 karyotype of this species. It will be informative to explore the origins of other species where n≠ 6, such as species in the genus *Diploscapter*. In *A. rhodensis*, these events may have been relatively recent, phylogenetically speaking, as there are still clear blocks of genes of different Nigon unit origin within the fusion chromosomes LG2, LG3 and LG4.

Fusions are not unique to *A. rhodensis*. NE is intact in *A. rhodensis* (Ar_LG1), *C. elegans* (Ce_V) and *H. contortus* (Hc_V) but has fused with NN (n1n2) in *P. pacificus* to form Pp_I. As noted previously [78], Pp_I is a fusion chromosome incorporating components of Ce_V (NE-derived) and Ce_X (NN-derived), but we note that the continued distinctiveness of the NE-derived and NN-derived components within Pp_I suggests that this fusion is phylogenetically recent. The two Nigon unit components within Pp_I retain an arms-and-centres long-range structure that is presumably derived from the original separate chromosomes, with high repeat density in the ancestral arms and high gene density in the ancestral centres. It has been proposed that the NE-NN fusion observed in *P. pacificus* is ancient, based on identification of a NE-NN like junction fragment in the genome sequence of the tylenchine (Clade IV) nematode *Bursaphelenchus xylophilus*, which is an outgroup to the rhabditine species. This apparent conservation of the junction fragment conflicts with the within-chromosome rearrangements dynamic observed elsewhere in the genome, and may be a chance homoplastic association of NE and NN elements in this species. In *O. tipulae*, NE has fused with NX to form Ot_X, and the NN unit forms a chromosome on its own (Ot_V).

The X chromosomes of the species analysed always contain the NX unit either as the sole component of the X (Pp_X and Ar_LG5X), or associated with other Nigon units: NN in Ce_X and Hc_X, and NE in Ot_X. The complex history of the NN, NX and NE units requires additional analyses, as it is unclear if NN belongs to NE (as found in Pp_I), to NX (as found in Ce_X and Hc_X) or if it is a unit of its own (as found in Ot_V). The number of chromosomally-assembled rhabditine genomes is still too few to fully define the ancestral gene content of Nigon units.

Previous studies have shown that segregation of the *A. rhodensis* X chromosome follows different programs depending on organismal sex and type of gametogenesis [17, 20, 21]. No recombination of the X is observed during hermaphrodite oogenesis and spermatogenesis leading to nullo-X oocytes and diplo-X sperm. Additionally, during outcrossing the X chromosome is always transmitted from father to son (males produce exclusively haplo-X sperm). Some predictions from this atypical inheritance are that the genes on the X will be more exposed to selection, genetic diversity on the X will be maintained for longer periods of time and essential genes will tend to migrate from the X to autosomes leading to a reduction in size. The X chromosome of *A. rhodensis* has many distinctive features. It is much smaller than the autosomes, representing only 6% of the genome and containing just over 600 genes. It carries no detected tRNA genes, and fewer conserved genes were found on the X compared to the autosomes. In *C. elegans* the X chromosome carries 44% of all tRNA genes and there is no marked exclusion of conserved genes [69]. In *A. rhodensis*, the X is inherited from father to son and is haploid in males, and thus genes on X chromosomes transmitted between males will be more exposed to natural selection, which will tend to exclude essential genes from the X [20]. In addition, the lack of recombination in hermaphrodites will slow down the removal of deleterious mutations.

Within-strain and between-strain genetic diversities were higher on the *A. rhodensis* X chromosome than on the autosomes. In *A. rhodensis* XX progeny resulting from a selfer always retain maternal heterozygosity on the X [20]. In our inbreeding, bottlenecking was performed by isolating a selfing hermaphrodite every few generations, and thus the X chromosome will only have recombined in females during the population expansion from each isolated hermaphrodite. As several generations occurred between each hermaphrodite isolation we expect that the X would become homozygous at a much slower rate than the autosomes.

Another fundamental question in *A. rhodensis* biology is the mechanism that controls female *versus* hermaphrodite sex determination in XX animals. Females and hermaphrodites are karyotypically identical, and are thought to be genetically identical. Our transcriptomic comparisons between female, hermaphrodite and converted female early larvae (L2) show that the sex differentiation process modulates the expression of many genes, with ~20% of all genes found differentially expressed between females (normal and converted females) and the hermaphrodites. We noted that *A. rhodensis* orthologues of some genes previously shown to be active in *C. elegans* sex determination were differentially expressed in females *versus* hermaphrodites. *A. rhodensis gld-1* was 200-fold overexpressed in females. In *C. elegans* hermaphrodites, *gld-1* is necessary for normal oogenesis and promotes spermatogenesis in hermaphrodites [79, 80]. *A. rhodensis tra-1* was 4-fold overexpressed in females. The zinc finger protein TRA-1 is the terminal regulator of the sex determination pathway in *C. elegans*, where it promotes female development in somatic tissues [81, 82]. Its role in sex determination is conserved in nematodes, making it an interesting target for functional studies [83]. We also identified a DM (*doublesex*/*mab-3*) domain transcription factor, homologous to *C. elegans dmd-10* and *dmd-11* that was 200-fold overexpressed in hermaphrodites. DM domain genes regulate sex determination and sexual differentiation processes in a number of organisms [84], but specific roles for *C. elegans dmd-10* and *dmd-11* have not yet been elucidated. RNAi knockdown of this locus in hermaphrodites resulted in the production of more female progeny, suggesting arole for this DM domain-coding gene in *A. rhodensis* sex determination.

The near identical expression pattern between converted females and females shows that the conversion of L1 hermaphrodite-fated larvae by exposure to DA is almost complete and that the initial sex decision can be overridden almost fully by environmental cues. The age and sex of an *A. rhodensis* mother affect the proportion of each sex in its progeny [19] and thus it is likely that maternal effects may directly establish the distinct developmental trajectories of females and hermaphrodites. However, the female *versus* hermaphrodite decision could also be modulated by environmental cues acting during embryogenesis and the L1 stage. These maternal and environmental effects could be modulations of what is essentially a random sex determination (RSD) system [85]. RSD occurs when fluctuations in the expression of genes at the top of the sex determining cascade or “developmental noise” are enough to canalise sexual fate down contrasting paths. An RSD component in *A. rhodensis* is plausible as females and hermaphrodites likely share the same genome, all sexual morphs are produced in a single environmental condition and the proportion of each sex produced varies greatly between mothers (although they are from inbred lines).

*Auanema* nematodes thus offer a fascinating and potentially highly informative model system for depth exploration of the origin of novel traits and their consequences. The genomic and transcriptomic resources we present for *A. rhodensis* will be critical for future analyses of the origins of new chromosomes in an otherwise stable karyotypic system, the biology of the highly regulated pattern of X chromosome segregation, the dynamics of mating system evolution, and the evolution of sex determination mechanisms. Towards this, we have identified the orthologs of 16 main sex determination genes, including *tra-1/2*, *gld-1*, *her-1* and *fem-1/2* (Supplementary Table 5), which are clear candidates to investigate the sex determination mechanisms in *A. rhodensis*. In parallel, we are developing reverse genetic and functional genomic technologies for these species, and these promise routes to rapid validation of hypotheses of gene function [16]. The *A. rhodensis* sex determination system may integrate genetic, maternal, environmental and random components, and this nexus of interacting components will also become amenable to manipulation and dissection. Genetic and genomic investigation of additional *Auanema* and closely related rhabditomorph species will contribute to a complete understanding of the origins and maintenance of this unusual mating system.

## Author Contributions

Conceptualization: ST, APS and MB; Generation of AILs, DNA and RNA extractions: MP; Genome assembly and annotation: GDK and ST; Genetic linkage map: ST; Genome analysis: ST, MB, APS; Transcriptomic analysis: ST; RNAi: SA, DC and ST; Data curation: ST and GDK; Supervising: APS and MB; Manuscript writing: ST, MB, APS and SA with input from all authors.

## Acknowledgments

The authors are very grateful to Stephen Doyle (from the Sanger Institute) for providing early access to the *Haemonchus contortus* genome and annotation, and to Fabrice Besnard for discussion about *O. tipulae* linkage groups. We thank staff of Edinburgh Genomics and UT Southwestern facilities for support in sequencing.

